# A larval zebrafish model of TBI: Optimizing the dose of neurotrauma for discovery of treatments and aetiology

**DOI:** 10.1101/2024.06.14.599067

**Authors:** Laszlo F. Locskai, Taylor Gill, Samantha A.W. Tan, Alexander H. Burton, Hadeel Alyenbaawi, Edward A. Burton, W. Ted Allison

## Abstract

Traumatic brain injuries (TBI) are diverse with heterogeneous injury pathologies, which creates challenges for the clinical treatment and prevention of secondary pathologies such as post-traumatic epilepsy and subsequent dementias. To develop pharmacological strategies that treat TBI and prevent complications, animal models must capture the spectrum of TBI severity to better understand pathophysiological events that occur during and after injury. To address such issues, we improved upon our recent larval zebrafish TBI paradigm emphasizing titrating to different injury levels. We observed coordination between an increase in injury level and clinically relevant injury phenotypes including post-traumatic seizures (PTS) and tau aggregation. This preclinical TBI model is simple to implement, allows dosing of injury levels to model diverse pathologies, and can be scaled to medium- or high-throughput screening.

## Introduction

Traumatic brain injuries (TBI) lead to diverse pathophysiologies such as tissue damage, blood flow abnormalities, metabolic disturbances, oxidative stress, and inflammation^1^. TBIs impact tens of millions of people worldwide, with estimates varying greatly due to differences in TBI classification and underreporting of non-hospitalized TBI patients^2,3^. TBI has devastating effects on children, domestic abuse victims, military personnel and athletes. TBI can result in tragic and long-term consequences such as disability, behavioural and emotional changes, seizures, and an increased risk of developing dementia/neurodegenerative diseases^4-9^. We recently established a simple, accessible method to deliver blast TBI to larval zebrafish^10^. The method employs dropping a weight on a closed syringe containing zebrafish in their typical aqueous media, to create blast pressure waves in the syringe (Figure 1)^10^. This TBI method offers various experimental and bioethical advantages, and we seek to improve upon and share those methods here.

**Figure 1.**
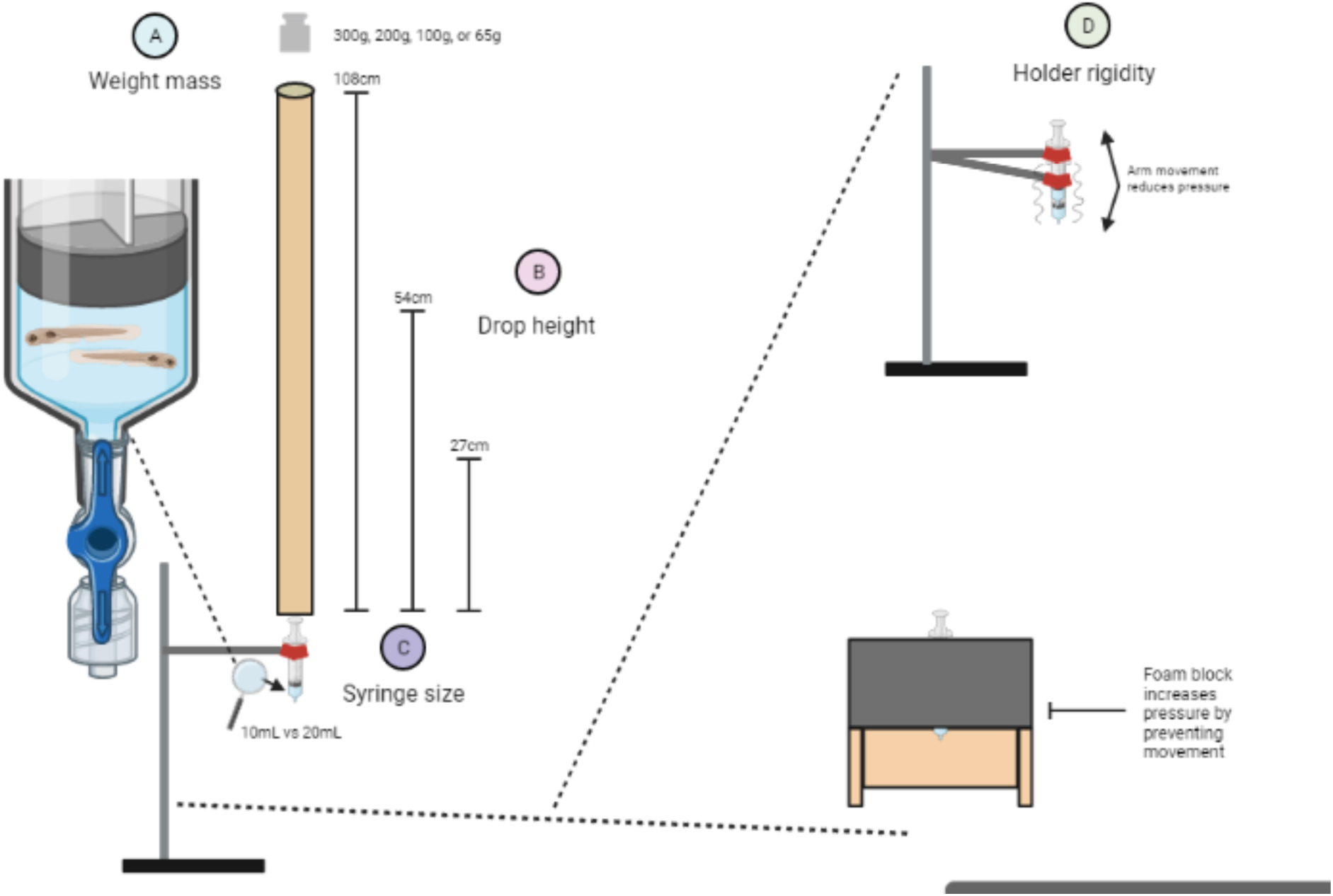
Larval zebrafish model of TBI and the modifiable aspects investigated herein. The larval zebrafish TBI model provides an elegantly simple system to create a blast pressure wave and induce TBI. A weight is dropped onto the plunger of a syringe housing dozens of zebrafish larvae in media. The plunger depresses when struck by the falling weight, and this transduces a pressure wave through the liquid media and thus through the zebrafish larvae. In the present study, we modified the following aspects of the larval zebrafish TBI model to increase the blast pressure wave created and allow the user to administer a variety of injury doses: **A**) The mass of the weight that was dropped. **B**) The height from which the weight was dropped. **C**) The size (volume) of the syringe that housed the group of zebrafish larvae. **D**) The rigidity of the syringe mount, where a syringe clamp holder had arm movement which dissipates pressure upon impact *vs.* a foam block syringe holder which halted syringe movement upon weight impact. Parameter changes are colour coded throughout the manuscript to match the numbered colours of this figure.

Preclinical TBI models across various species and modes of injury have been productive in advancing our understanding and treatment of neurotrauma. Each model has benefits and challenges. The most prominent research uses rodent TBI models via fluid percussion injury, controlled cortical impact injury, weight-drop impact injury, or blast injury^11-14^. Large animal micropig TBI models provide valuable information about injury events, such as biomechanical aspects of injury that cannot be recreated in small animal models^15-17^. *Drosophila* TBI models allow cell-specific genetic manipulations ideal for dissecting injury pathways^18,19^. While these animal models are valuable, they are accompanied by various limitations that warrant consideration. For example, mammalian models are not readily accessible to optogenetic measures or manipulations, especially *during* physical injury, and this has hampered an appreciation of injury-relevant neural activity, neurotransmitters and receptors. Neurotrauma models in *Drosophila* offer increasing innovations and high throughput, but the drug screening is laborious and the central nervous system (CNS) biology of *Drosophila* and TBI patients is disparate (e.g. roles for glia in synapse homeostasis, glymphatic system for managing proteostasis, etc.) and requires care in translating the outcomes.

Several adult zebrafish TBI models exist such as stabbed lesion brain injury, closed head pulsed ultrasound injury, blunt force TBI, and laser irradiation TBI^20,21^. In larval zebrafish, TBI has been modelled by stabbed lesion brain injury, linear deceleration induced mild TBI, and the blast pressure TBI model discussed in this study^10,22-25^. Adult and larval zebrafish models both have unique strengths and both support translation to adult humans, but here we focus on the benefits of larval zebrafish models.

Zebrafish larvae are a powerful animal model of human disease because they have complex physiological systems akin to humans, making them advantageous compared to in vitro or invertebrate models. The intricate but accessible CNS of zebrafish larvae enables models of neurological diseases such as epilepsy, Parkinson’s disease, and tauopathies^10,26-31^. Conserved proteins and biochemistry ^32,33^ enable modelling of neurodegeneration and dissection of pathomechanisms^34-37^. Larvae can be obtained in high quantities (hundreds per breeding tank) meaning thousands of samples can be run in a variety of assays each week. Zebrafish larvae are small, being several millimeters in length (and growing rapidly), making them amenable to 96-well plate formats for analysis. Zebrafish larvae are amenable to high throughput assays^38,39^, which is a powerful untapped opportunity for TBI therapeutic research with the potential of testing small molecule libraries, CRISPR screening of genetic targets, and manipulation of molecular pathways to dissect pathophysiological events. High throughput larval zebrafish models are ideal for discovery of seizure/epilepsy therapeutics, which represent a major target in the form of post-traumatic seizures (PTS) and post-traumatic-epilepsy (PTE). For example, clemizole is in phase II clinical trials for Dravet’s syndrome, a genetic epilepsy, which was discovered through assessing a larval zebrafish disease model with high throughput assays^40^. Clemizole and other zebrafish-based drug discoveries highlight the translational capabilities and practicality of larval zebrafish models for the discovery of clinical therapeutics^33,41-47^. Neural activity and other events can also be modelled *during* injurious impact using larval zebrafish, due to their transparent nature and innovation with calcium imaging paradigms like CaMPARI^10,48-50^.

Limitations of larval zebrafish for neurotrauma research include the paucity of robust assays of cognition (learning and memory), a lower husbandry temperature (28°C) compared to human physiology, and a relatively modest toolkit for cell- or tissue-specific genetic modifications (compared to *Drosophila* or mouse). These are offset by additional benefits such as diurnal habit (contrasting nocturnal rodents and associated logistics of investigating sleep homeostasis, behavioural metrics and glymphatic clearance, etc.). Zebrafish also provide important bioethical considerations within the principles of The Three Rs: Replacement, Reduction and Refinement^51^. In our model, zebrafish larvae can serve to Replace adult and mammalian animals that would otherwise need to be exposed to neurotraumatic interventions.

We have previously demonstrated that our TBI model produces TBI pathologies by examining dozens of individual larvae after TBI, with individual variability and robust statistics available^10^. These pathologies can be investigated in short timeframes compared to rodent models. We found that neural activity increased during TBI using optogenetics and that PTS and reduced blood flow occurred within hours post-TBI. Neuronal cell death and progressive tau pathology were apparent days after TBI. Excitingly, post-TBI therapeutics were tested using our model and we discovered treating post-TBI neurovolatility (aberrant neural activity such as clinical/non-clinical seizures)^52^ with anti-epileptic drugs reduced dementia pathology, whereas increasing neurovolatility with convulsants worsened pathology^10^. Our finding that therapeutically applying the anti-convulsant retigabine prevents TBI-mediated dementia pathology^10^ was validated by others using blast TBI in rodents^53^. Surprisingly, our data revealed that one convulsant was unexpectedly protective due to off-target effects which would not have been discoverable *in vitro* ^10^.

Here we seek to expand upon the model and make the methods openly accessible. Our design principles for the optimization of the larval zebrafish TBI model include: *i*) Retaining the method’s elegant simplicity, in that the materials required are inexpensive and accessible in most labs, and there are no intellectual property constraints. Any zebrafish researcher can access this TBI model. *ii*) TBI levels are optimized to trade-off between maximizing injury intensity and larval survival. *iii*) Various injury intensities can be implemented with reliable consistency to provide a low-cost dose-response paradigm. *iv*) Animal ethics are paramount, where larval zebrafish fulfil Replacement principles^51^ for adult rodent TBI models; Moreover, we present methods allowing researchers to Refine^51^ the method using an inexpensive pressure transducer rather than trial-and-error on animals.

We have empirically adjusted components of our TBI method to characterize the pressure dynamics and applied this information to optimize injury for the study of moderate-to-severe TBI. The complex heterogenous nature of TBI is a primary factor for the lack of successful translation of therapeutics in clinical settings^54,55^. By optimizing our parameters, different TBI intensities can be modeled in our system to further study heterogenous injury pathomechanisms and potential therapeutics. A contemporaneous publication provides step-by-step instructions with set-up and trouble-shooting advice^56^, and together these writings are intended to allow easy implementation of zebrafish TBI research in any Institution. Various TBI model parameters were modified (see Figure 1) including the mass of the dropped weight, the height the weight was dropped from, the size of the syringe housing the larvae, the rigidity of the syringe holder, and the number of times the weight was dropped per injury group. We chose the more ethically favourable route of replacing animals with a custom pressure sensor to optimize pressure levels. Once optimized mechanically, we then validated the outcomes against the biological effects of various injury severity levels, including by measuring PTS, stimuli response, and tau pathology, representing both behavioural and neuroanatomical outcomes of neurotrauma.

## Methods

### Animal ethics

Zebrafish were raised and maintained following protocol AUP00000077 approved by the Animal Care and Use Committee: Biosciences at the University of Alberta, operating under the guidelines of the Canadian Council of Animal Care. The fish were raised and maintained within the University of Alberta fish facility under a 14:10 light: dark cycle at 28°C as previously described^57^.

### Larval zebrafish TBI model

In the present study, we refined our previously described larval zebrafish blast pressure TBI model^10^ (Figure 1). Detailed step-by-step instructions and setup considerations have recently been made available^56^. Larvae were placed into a 10mL or 20mL syringe (Becton Dickinson #309604, #302830) with 1mL of E3 media, which was then closed with a Luer-Lok stopper valve attachment (Cole-Parmer #UZ-30600-00). The syringe was placed in either: *1)* a three-prong clamp centred on a support rod and stand (E.g. Fisher # S41710) or *2)* a more rigid foam block with a hole drilled into the center matching the syringe size, fashioned from a standard yoga foam block (23 x 15 x 7.6cm dimensions). Once mounted the syringe plunger was centered underneath a guide tube made from rolled paper with a circumference slightly bigger than the largest weight used (40mm diameter for a standard 38mm calibration weight; tube heights measured 27cm, 54cm, or 108cm). A weight (e.g. scale calibration weight) was dropped into the guide tube, creating a pressure wave upon impact. Between weight drops, the syringe stopper was opened to remove air bubbles and reset larvae back into the syringe barrel, and the time between drops was <10 minutes.

### Pressure detection setup

To measure pressure in the syringe, we used an Arduino Uno Rev3 microcontroller board (Arduino #A000066) and an automotive fuel line pressure transducer (AUTEX GSND-0556629788) as described previously^56^. The Arduino board was connected in a circuit to the pressure transducer and a photoresistor (eBoot EBOOT-RESISTOR-05) using a breadboard (Haraqi ESH-PB-01). The photoresistor was positioned opposite from a light source at the top of the guide tube. When the weight was dropped into the guide tube the plane of light would break, altering the photoresistor voltage and triggering the recording. The pressure transducer then measured for one second after the photoresistor voltage dropped (alternatively the light was kept off if longer recordings were required). The pressure transducer output was collected in the Arduino software (Arduino IDE 2.0.3) with a setting of 2,000,000 baud. The output was converted to pressure (kPa) with the formulas [voltage = (5 * output)/1023] where output is the values given by the pressure transducer, [PSI = (voltage – 0.5) * (37.5), and [kPa = PSI * 6.895]. For pressure wave measurements, the first pressure spike reading from baseline until the first measurement back at baseline was considered a single pressure wave.

To validate the transducer calibration, static weights of varying mass were rested on the syringe plunger, and readings of the transducer were compared to expected values. Expected pressure values of each weight at rest are calculated using Pascal’s principle: pressure = weight / cross-sectional area of the syringe barrel. Caution may be warranted that our calibration of the pressure used a static weight, and information regarding the pressure transducers’ dynamic performance is not provided herein.

### Behavioural detection of post-traumatic (PTS) seizures

To measure PTS, we used a locomotor assay previously described^10,49,50,58^. In short, larval zebrafish were subjected to TBI at 6 days post fertilization (dpf) using the methods described. Larvae were used at 6dpf because their CNS is well-developed by that age, the larvae are robust to manipulation, and it was convenient to our workflow, but other ages could be useful for other workers. Larvae were then placed into a 96-well plate in 200uL of E3 media and recorded for 30 minutes using the behavioural software EthoVisonXT (11.5) 45 minutes after TBI.

### Manual scoring of seizures

Seizures were assessed manually to account for the confounding effect of larval inactivity at higher injury intensities. Larvae were administered TBI of varying levels and then recorded for 2 hours using EthovisionXT as described above. A high activity bout threshold was set at 64% of the highest maximal activity movement detected in the uninjured group. This threshold detected abnormally high bouts of activity and when they occurred. The time points of these high-activity bouts were then scored (by blinded observers) for the presence of seizure characteristics. Larval zebrafish seizures are defined by three stages, where stage I seizures are increased activity, stage II seizures are whirl-pool motions, and stage III seizures are clonic seizure bouts with periods of loss of posture (LOP) and sinking^59,60^. Here, stage I seizures were scored as distinct hyperactive movements with slight convulsive behaviour and minor LOP (LOP <1 second/larvae still moving). Stage II/III seizures were scored as whirlpool movements, clonic seizure events, and LOP (fully on side with inactivity >1 second). Larvae that floated in their well and did not display any movement were considered dead and removed from the data.

### Larval zebrafish fin poke response

To further assess larval inactivity after TBI, the stimuli response of larvae was assessed post-TBI using a fin poke response assay. This assay is used to measure loss of consciousness after anesthetization in zebrafish^61^. Larvae at 6 dpf were injured with various TBI intensities and placed in a 10mL petri dish by group. The larvae were poked once with a sharp metal point on the posterior fin and observed for response every 5 minutes for 30 minutes (blinded). Responsive larvae immediately swim away, whereas unresponsive larvae do not respond. The heartbeats of the larvae were assessed after quantification to ensure no dead larvae were included in the data.

### Tau-GFP quantification

To measure the formation of GFP+ tau puncta we used *Tg(eno2:Hsa.MAPT_Q244-E372−EGFP)ua3171* larvae (ZFIN ID: ZDB-ALT-211005-6) crossed with *Tg(eno2:hsa.MAPT-ires-egfp)Pt406* larvae (ZFIN ID: ZDB-ALT-080122–6), which express full-length 4R human tau and the 4R domain of human tau linked to GFP, both under the eno2 promoter^10,62^. These fish have been previously validated to develop tau GFP+ aggregates after TBI^10^. Larvae subjected to TBI (as described above) at 3 dpf were anesthetized at 7 dpf with MS-222 and imaged under a fluorescent dissecting scope (Leica M165 FC). GFP+ puncta were counted from a lateral view (blinded).

### Statistics

All statistical analyses were done using GraphPad Prism Software (version 10.0.0). Mean and maximum pressures were calculated +/- standard error from the mean in units of kilopascals (kPa). Relative standard deviations were calculated taking the proportion of the SD to the mean, multiplied by 100. One-way ordinary ANOVA tests were used for statistical analysis of EthovisionXT seizure data followed by Dunnett’s multiple comparison of means. Linear relationships were analyzed using simple linear regression, whereas binary relationships were analyzed with multiple or simple logistic regression.

## Results

### Characterizing pressure for the optimization of injury

Each of the four adjustable parameters (denoted as “A” thru “D” in Fig. 1) was combined to provide a broad range of TBI pressures that span three orders of magnitude (Fig. 2). The weight drops created a pressure waveform consisting of multiple pressure waves as the weight bounced (Fig.2A,B), so we first focused on the maximal pressure of each parameter change individually. Changes to the mass of the weight dropped (as depicted in Fig. 1A) had the largest impact on maximal pressure ranging from a 6- to 21-fold increase across all TBI setups, whereas the drop height of the weight (Fig. 1B) had the second largest effect increasing maximal pressure up to >3-fold (Fig 2C-F, Supplemental Tables S1-S4).

**Figure 2.**
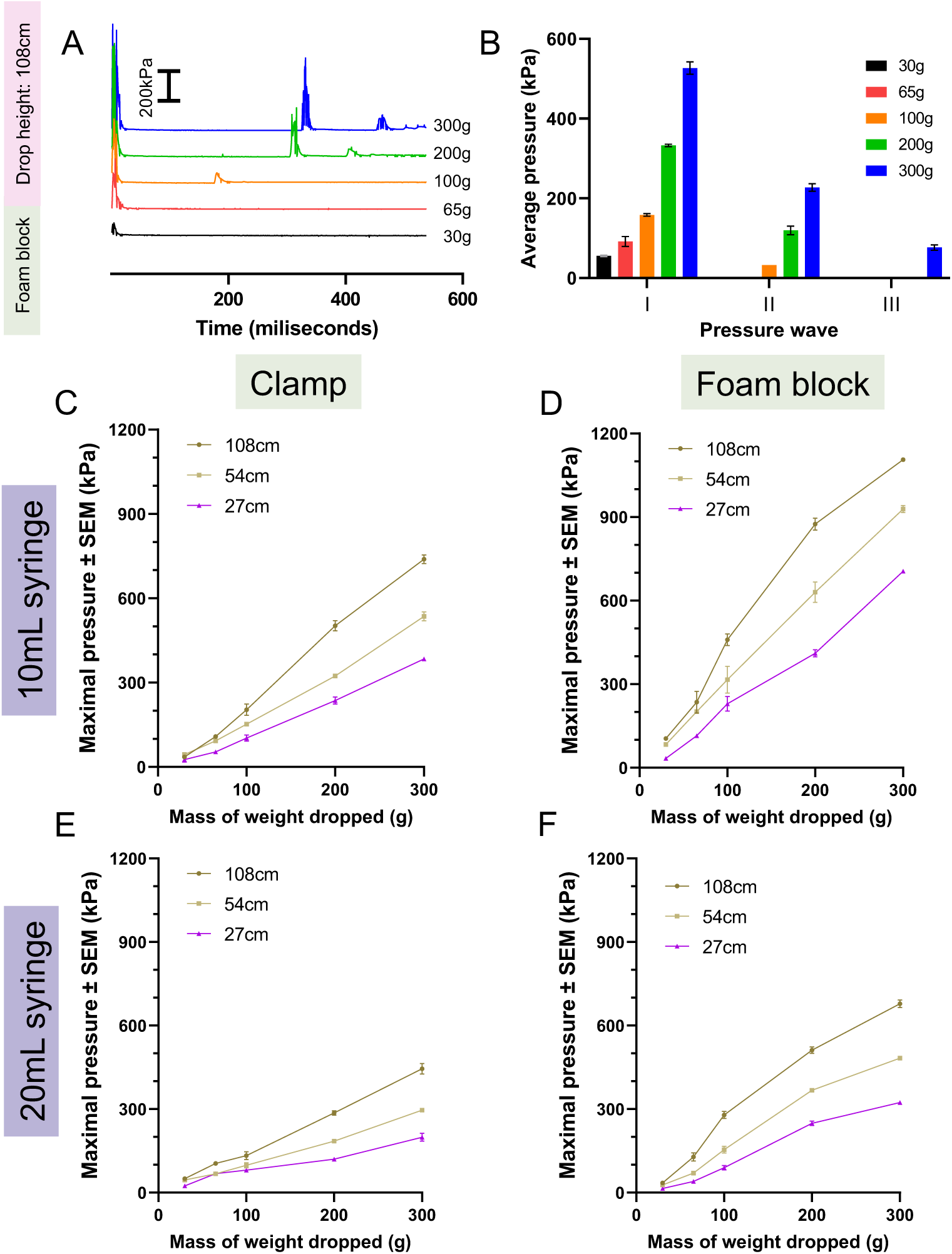
Maximal pressure created in alternate configurations of the larval zebrafish TBI model. To optimize TBI forces, four equipment parameters were adjusted in concert: *(i)* A 10mL or 20mL syringe was used, and *(ii)* the syringe was held by either a clamp or a rigid foam block. Next, *(iii)* weights of various masses were dropped *(iv)* from various heights (108cm, 54cm, 27cm). Weights ranging from 65-300g were dropped onto the syringe, and the pressure levels generated for each configuration were measured using Arduino IDE software. Pressure values displayed are the mean values of the maximal pressure reading achieved from three independent weight drop recordings. **A, B**) Example time course of pressure dynamics and average pressure generated per pressure wave in the 108cm/20mL syringe/foam block TBI setup. This data is duplicated & expanded in Sup. Figs. 2-5 to highlight that additional pressure waves are generated due to the weight bouncing on impact. **C-F**) Maximal pressure measurements for each indicated parameter combination. Maximal pressure can be manipulated across three orders of magnitude through changing various TBI assay settings. Alterations to the mass of the weight dropped had the largest impact on maximal pressure, followed by changes to the drop height of the weight. Using a foam block to hold the syringe and decreasing the syringe size both modestly increased maximal pressure.

Decreasing syringe size when using the clamp syringe holder (Fig. 1C) decreased maximal pressure up to 0.7-fold for lower drop weight masses but modestly increased (2-fold) maximal pressure at higher drop weight masses (Fig. 2C.E, Supplemental Table S1 & S2). Decreasing syringe size when using the foam block syringe holder resulted in a 1.62- to 3.05-fold increase in maximal pressure (Fig. 2D,F, Supplemental Table S3 & S4). Increasing syringe holder rigidity by using the foam block syringe holder instead of the clamp (Fig. 1D) increased maximal pressure 1.29- to 2.98-fold when using a 10mL syringe (Fig. 2C,D, Supplemental Table S1 & S3). Comparatively, the rigid foam block syringe holder decreased maximal pressure up to 0.59-fold for 30g (all heights) and 65g at 27cm height but increased pressure up to 2.10-fold otherwise when using a 20mL syringe (Fig. 2E,F, Supplemental Table S2 & S4). Overall, maximal pressure was achieved by using the 10mL syringe, rigid foam block holder, and 300g weight dropped from 108cm.

Consistency of pressure values across parameter changes was compared using the relative standard deviation (RSD) of the maximal pressure (Sup. Fig. 1). RSD decreased as the mass of the weight increased, except for the 65g weight which displayed a lower RSD versus the 30g and 100g weight (Sup. Fig. 1). The lower RSD values for 65g weights may be because it had a diameter similar to the higher weights, whereas the 35g and 100g weights were smaller. The diameter of the weight being mismatched to the diameter of the guide tube used may introduce variability when dropped, perhaps by being more prone to falling at different angles.

Although <u>maximal</u> pressure (discussed above) is a useful metric to compare pressure between TBI setups, secondary pressure waves generated add complexity to the total injurious force created (Fig. 2A,B). We measured the <u>average</u> pressure per wave generated after impact to investigate how parameter changes impact secondary pressure waves. Overall, an increased mass of the weight dropped and decreased syringe size increased the number of additional pressure waves generated (Sup. Fig. 2-5, Supplemental Tables S1-S4).

These secondary pressure waves add complexity when comparing similar maximal pressure measurements (as calculated in Figure 2) of different TBI setups. For example, 200g dropped on the 10mL syringe/foam block generated a primary and secondary pressure wave 210% and 75.4% of the pressure of the initial wave generated by 100g in the same setup, respectively (Sup. Fig. 4B, Supplemental Table S3). The injurious impact of 200g would be more than double the impact of 100g, followed by a second impact on par with the primary impact from 100g, meaning the pressure experienced by larvae is much greater than that suggested by maximal pressure or the primary waves alone. We suggest that secondary pressure waves should be considered when cross-comparing the injury phenotypes measured in each TBI model setup at similar pressure levels.

### Post-traumatic seizures (PTS) increase with injury severity, but severe injury confounds behavioural tracking of seizures

To assess PTS, we used a well-established behavioural assay that quantifies larval movement, using the software EthovisionXT, as a proxy of seizure intensity following TBI. Our goal was to measure how changing our TBI model parameters impacts the pathobiological outputs of neurotrauma injury and how these pathobiological changes are related to pressure measurements.

First, we measured the locomotor activity of larvae injured in the 20mL syringe and clamp TBI setup, which we have previously used to detect seizures^10^. Overall, TBI increased locomotor activity as the height of the drop weight was increased up to more than 3-fold, but activity decreased when injury became too high (Fig.3A). Larvae injured with the 100g and 200g weight showed hyperactivity versus no TBI larvae as weight drop height increased to 54cm and 108cm (Fig.3B,C, p<0.05-0.0001; effect sizes show doubling or tripling of activity levels and varying with mass of the weight and height). Larvae injured by the 300g showed hyperactivity at 27cm and 54cm weight drop heights, but activity decreased for the 108cm injury group (Fig.3D, p<0.05).

**Figure 3.**
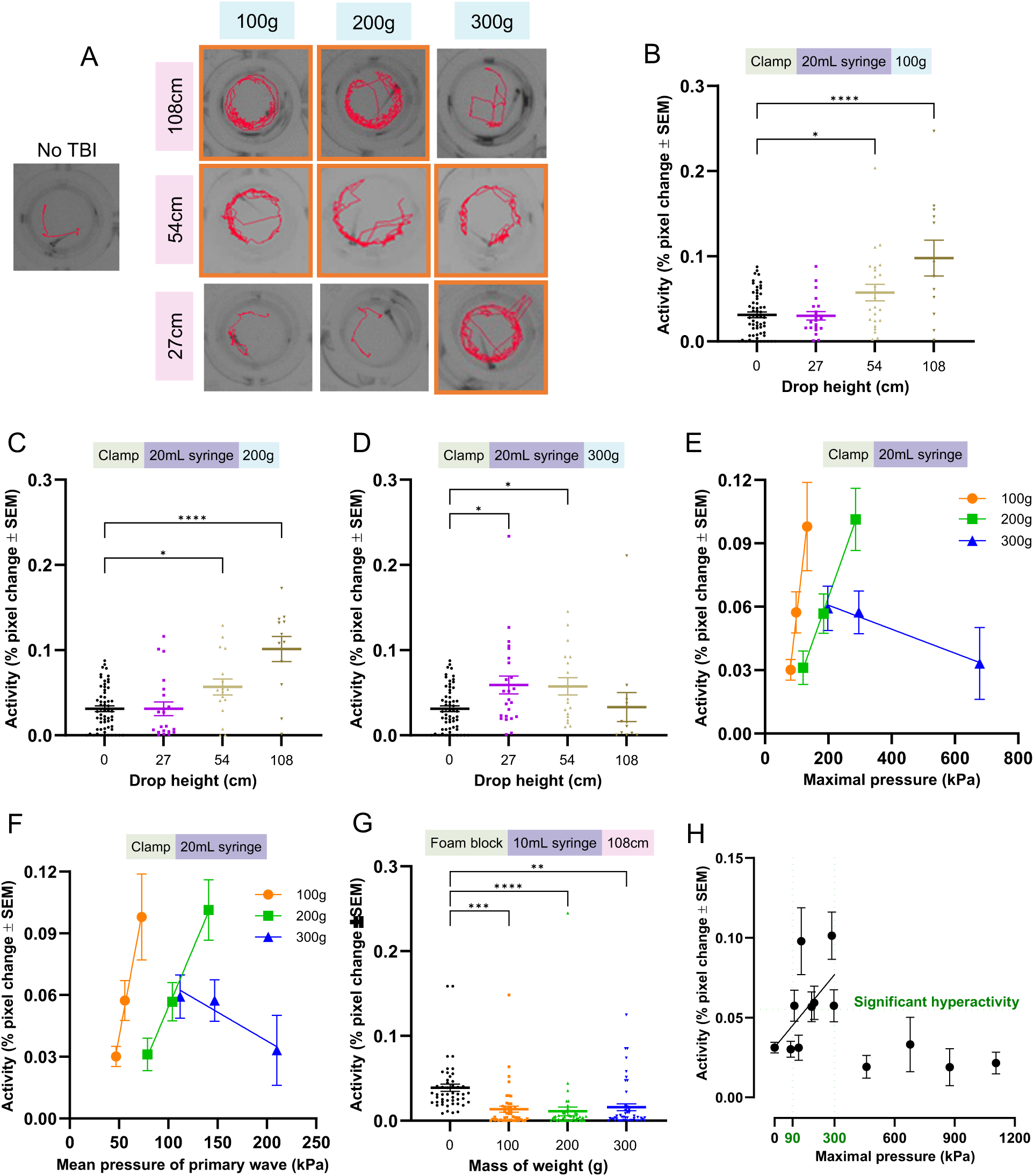
Seizure-like locomotor activity of 6 dpf larvae following TBI increases at moderate injury levels but decreases at more severe injury levels. Following TBI injury with various parameters, seizure-like activity was assessed by placing the larvae in a 96-well plate recorded using EthovisionXT. Each data point represents an individual zebrafish larva. **A**) Exemplar activity traces of individual zebrafish larvae within wells of a 96-well plate. The red depicts the movement pattern of the larvae during one minute of activity and orange boxes denote groups that displayed significantly increased mean activity. **B-D**) Mean locomotor activity of larval zebrafish after dropping 100g (B), 200g (C), or 300g (D) from various heights onto a 20mL syringe held by a clamp compared to a no TBI control. Each experimental group had two to four replicates. Each dot represents behaviour of an individual larva. No TBI control data (N = 60) is shared between each graph to prevent bias from splitting the individual control replicates amongst each comparative graph (B) 100g N= 21, 23, 12 for the 27cm, 54cm, and 108cm drop height groups;. C) 200g N= 21, 17, and 12 for the 27cm, 54cm, and 108cm drop height groups; D) 300g N= 24, 18, and 12 for the 27cm, 54cm, and 108cm drop height groups). **E, F**) Linear regression analysis of larval activity as a function of maximal pressure and the average pressure of the primary pressure wave, respectively, for each weight (maximal pressure R^2^: 0.99 p<0.01 all weights; average pressure of primary pressure wave R^2^: 0.99 (100 & 200g) and 0.90 (300g), p<0.05 (100g & 200g). **G**) Higher pressure foam block holder induced a decrease in zebrafish locomotor activity after dropping 100g (N= 48), 200g (N = 48), or 300g (N = 48) from 108cm onto a 10mL syringe held by a foam block compared to a no TBI control (N = 48) (four experimental replicates). **H**) Zebrafish locomotor activity as a function of maximal pressure experienced during injury shows a range of 90-300kPa where behavioural seizures are detected (green) and trend with increased pressure (line), whereas higher pressure induces decreased locomotor activity. A one-way ANOVA with Dunnett’s multiple comparisons of means was used to test for statistical significance for behavioural experiments (* = P<0.05, ** = P<0.01).

The 200g weight dropped from 54cm vs. 300g dropped from 27cm created similar pressure dynamics (Supplemental Table S2, sup. Fig. 3D,F) and both resulted in similar magnitude post-injury hyperactivity. 200g dropped from 108cm and 300g dropped from 54cm also produced similar pressure dynamics (Supplemental Table S2, sup. Fig. 3B,D), but the magnitude of hyperactivity for 300g/54cm injured larvae was 43.2% lower. Similarly, the pressure dynamics of 100g dropped from 108cm and 200g dropped from 27cm were comparable (Supplemental Table S2, sup. Fig. 3B,F), but only the 100g/108cm injured fish displayed seizure phenotypes.

Slight differences in pressure wave dynamics between TBI settings could contribute to larval injury. For example, the number of pressure waves differs for 100g dropped from 108cm and 200g dropped from 27cm (Sup. Fig. 3B,F). Additionally, TBI settings that caused a lower seizure phenotype compared to settings of similar pressure all had a shorter duration of time between the first and second pressure wave (average of 73-86 milliseconds), whereas TBI settings that had similar pressure values and seizure phenotypes only differed by 17 milliseconds on average. These data suggest that small differences between the pressure dynamics may alter phenotypes. Furthermore, the pressure values created by 200g from 27cm and 100g from 108cm are similar, but the gravitational potential energy of the former is 50% of the latter. Mechanical properties such as syringe and syringe holder movement etc. could potentially impact the injury and resulting phenotypes detected.

Overall, our larval TBI model was consistent across comparable groups and showed an upward trend in seizure activity as the drop height was increased for a given weight. Increases to maximal pressure and average primary wave pressure significantly correlate with the magnitude of detected seizures (Fig.3E,H,F, p<0.05; R^2^ = 0.99). These results suggest that the highest drop height is best for producing injury phenotypes with the largest dynamic range. When comparing TBI assay changes modifying one parameter at a time (e.g. drop height or weight) may minimize variables impacting pressure dynamics and produce the most consistent injury phenotype “dose response”.

Locomotor activity decreased at the highest injury level (Fig.3D), consistent with reduced locomotion TBI symptoms reported in human and animal models^21,23,63^. In one study consistent with our results, moderately injured rodents had higher locomotor activity, but severely injured rodents had decreased locomotor activity^64^. Additionally, intense larval zebrafish stage III seizures cause convulsive events and loss of posture (LOP) instead of regular swimming patterns, reducing detectable activity. To test if further increases in pressure and injury resulted in decreased locomotor activity, we administered TBI through our 10mL syringe/foam block setup (Supplemental Tables S1 & S3). All weight groups had significantly reduced activity (Fig. 3G, p<0.01-0.001). Seizure phenotypes were detectable for injuries of approximately 90-300kPa maximal pressure (Fig.3H, green), with additional factors aside from maximal pressure influencing the intensity of seizures as discussed above. injuries above 300kPa maximal pressure reduced activity (Fig.3H). These results support that when injury is increased above a certain level larval zebrafish display a reduced locomotor activity phenotype which may confound locomotor-based seizure detection.

To investigate PTS at higher injury levels, we manually quantified seizures. Larvae were injured in the 20mL syringe clamp or foam block TBI setup (Fig.4A). Stage I seizures were measured as hyperactive movements with minor convulsions and LOP, while stage II/III seizures were measured as distinct whirlpool motions and major convulsive episodes with distinct bouts of LOP^59,60^. The proportion of seizing larvae increased as the mass of the weight used for injury increased for the clamp syringe holder group (Fig. 4A). Larvae injured using the foam block syringe holder displayed a similar proportion of seizing larvae across all weight groups, but more high-intensity seizures were detected (Fig. 4A). Logistic regression analysis indicated the odds of a larva displaying seizure behaviour increased 0.5% for each 1kPa increase in maximal pressure or 64.7% per 100kPa (OR per kPa: 1.005, Table 1). Larvae in the clamp/200g vs foam block/100g injury groups experienced similar maximal pressures but displayed different phenotypes (Fig.4A,B), further highlighting that the number of pressure waves and timing between weight bounces may alter the injury phenotype.

**Figure 4.**
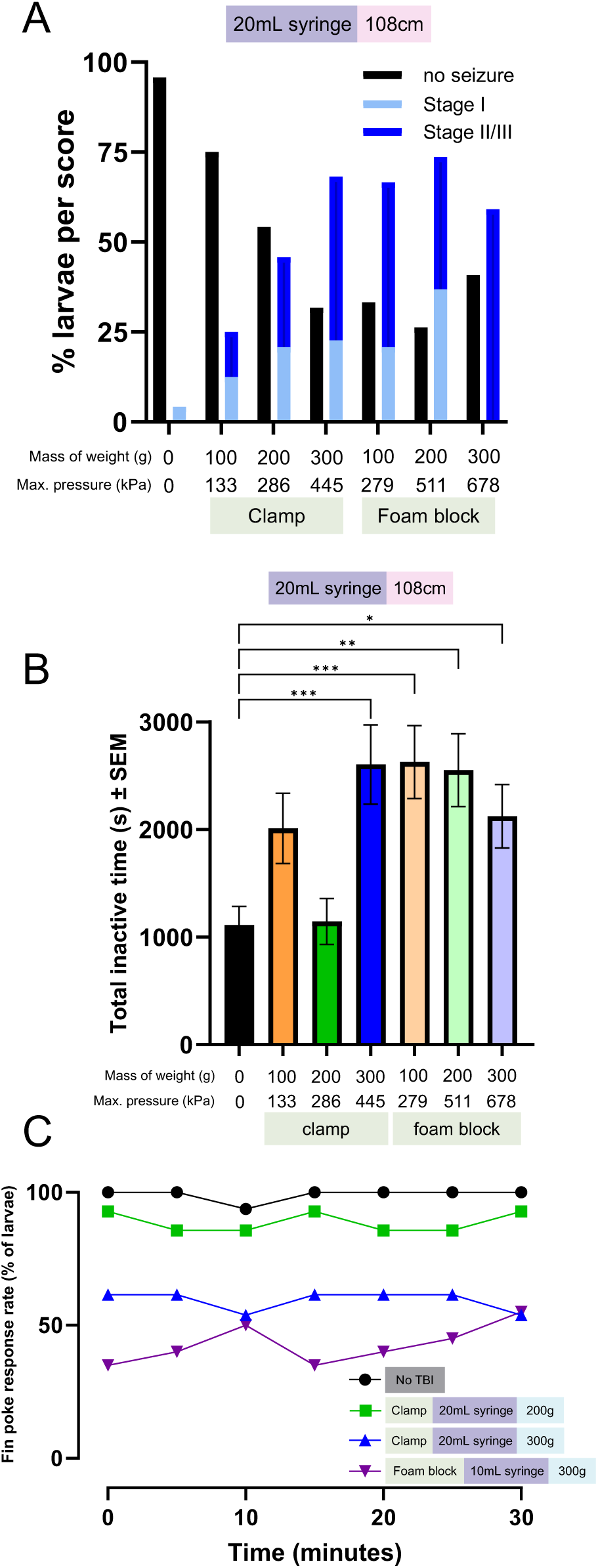
Post-traumatic seizures and unresponsive zebrafish behaviours increase with more severe injury. To assess decreased locomotor activity following higher injury and pressure intensities, seizures and loss of consciousness were measured manually. **A**) Zebrafish larvae were recorded after injury using either the foam block or clamp syringe holder. Larvae were manually scored (blinded) for the presence of seizure behaviour using three categories: no seizure, stage I seizure, or stage II/III seizure. The proportion of larvae displaying seizure trended with increased maximal pressure levels (N = 48 no TBI, N = 24 100g clamp, N = 24 200g clamp, N = 24 300g clamp, N = 24 100g foam block, N = 19 200g foam block, and N =22 300g foam block injured larvae. Each experimental group was repeated twice). **B**) A threshold of one minute or more of inactivity was set to quantify larval inactivity after injury in the videos used for manual seizure scoring. The total duration of these inactive bouts was quantified. Inactivity was significantly higher than no TBI controls for the highest pressure clamp injury group (300g) and all foam block injury groups (p<0.05-0.001). **C**) Following TBI larval zebrafish display a stunned phenotype where they do not respond to stimuli in the form of a fin poke, whereas non-injured fish respond vigorously. The proportion of larvae that exhibit this stunned phenotype increased for higher injury intensity groups versus lower injury groups (purple = highest intensity, N= 20; blue = middle intensity, N = 13; green = lowest intensity, N = 11. Each group had two replicates). Across experiments, dead larvae were removed from quantification. Larvae were verified as dead by a lack of heartbeat in the fin poke response assay and zero movements while floating during video recording. A one-way ANOVA with Dunnett’s multiple comparisons of means was used to test for statistical significance between injured larva inactivity and no TBI inactivity (* = p<0.05, **= p<0.01, *** = p<0.001).

**Table 1.** Logistic regression analysis considering how changes in TBI parameters impact the probability of larval zebrafish displaying TBI phenotypes after injury.

| TBI phenotype | Parameter change | Odds-ratio (β1) | 95% confidence interval | P value | Model fit (Area under ROC curve) |
| --- | --- | --- | --- | --- | --- |
| Seizures | Maximal pressure | 1.005 (per kPa) | 1.003-1.006 | <0.0001 | 0.77 |
| Stimuli response (30 minutes post-TBI) | Maximal pressure | 0.9978 (per kPa) | 0.9963-0.9992 | <0.01 | 0.79 |
| Tau pathology | Number of weight drops per injury group | 1.099 (per weight drop increase) | 1.013-1.194 | <0.05 | 0.81 |
| Tau pathology | Maximal pressure | 1.004 (per kPa) | 1.003-1.006 | <0.0001 | 0.81 |

Larvae with more intense seizures appeared to have long bouts of LOP and inactivity, so we next quantified cumulative inactive time across injury groups. Larvae injured in the clamp syringe holder had a non-significant increase in inactive time when injured with 100g, inactive time comparable to controls when injured with 200g, and a significant increase in inactive time when injured with 300g (Fig.4B, p<0.001). All larval groups injured in the foam block syringe holder displayed a significant increase in inactive time versus controls (Fig.4B, p<0.05-0.001). These results suggest that as injury intensity increases larval activity decreases even when seizures are apparent, further supporting our behavioural seizure results (Fig.3D,G).

To further characterize inactivity post-TBI we measured larval fin poke response, an assay used to assess larval loss of consciousness after anesthetization ^61^. The average stimuli response rate of larvae 30 minutes post-TBI decreased as the injury intensity increased (Fig.4C). Logistic regression analysis of the larval response rate 30 minutes post-TBI indicated a 0.22% decrease in stimuli response odds per 1kPa increase in maximal pressure, or 19% per 100kPa (OR per 1kPa: 0.9978, Table 1). These results suggest that more larvae develop a stunned/unresponsive injury phenotype as injury increases.

These data show that our larval zebrafish TBI assay can be optimized for the detection of PTS at multiple intensities. Altering weight and height parameters impacts behavioural seizure activity between 90-300kPa, but additional factors influence the magnitude of seizures. To reduce variability across TBI assay settings we recommend changing one parameter at a time. In our experiments, height was associated with increased seizure activity when weight was held constant, and a weight drop height of 108cm induced the strongest phenotype. We recommend a weight drop height of 108cm and adjustments to the weight if injury dosing is needed. Since increased injury suppresses activity, other assays like manual seizure scoring, optogenetic assays like CaMPARI, which we have validated during TBI^10^, or EEG may be needed for proper quantification.

### Tau aggregation increases with injury levels

Longer-term consequences of TBI include CTE, and a risk for AD. Thus, we next sought to identify the TBI parameters that produce a useful dynamic range for assessing tauopathy, using a genetically encoded Tau-GFP biosensor. This biosensor approach has been fruitful for assessing tauopathy in vitro^65,66^ and was recently deployed in larval zebrafish^10^. Tauopathy is scored in the spinal cord because it is easier to visualize and accurately quantify compared to tauopathy in the brain, and previous work showed the tau abundance in the two structures vary in good coordination^10^. To determine if increasing injury produces more tauopathy, various parameters were compared, while injury dose was further modulated by increasing the number of times the weight was dropped (Figure 5). Larval survival sharply declined in the highest-pressure TBI assay so survival was tracked from injury to quantification to highlight the tradeoff of increasing weight drops. Larval survival when injured in the clamp syringe holder was consistent with uninjured controls but starkly decreased when the number of weight drops was increased in the 10mL foam block setup (Fig.5E).

**Figure 5.**
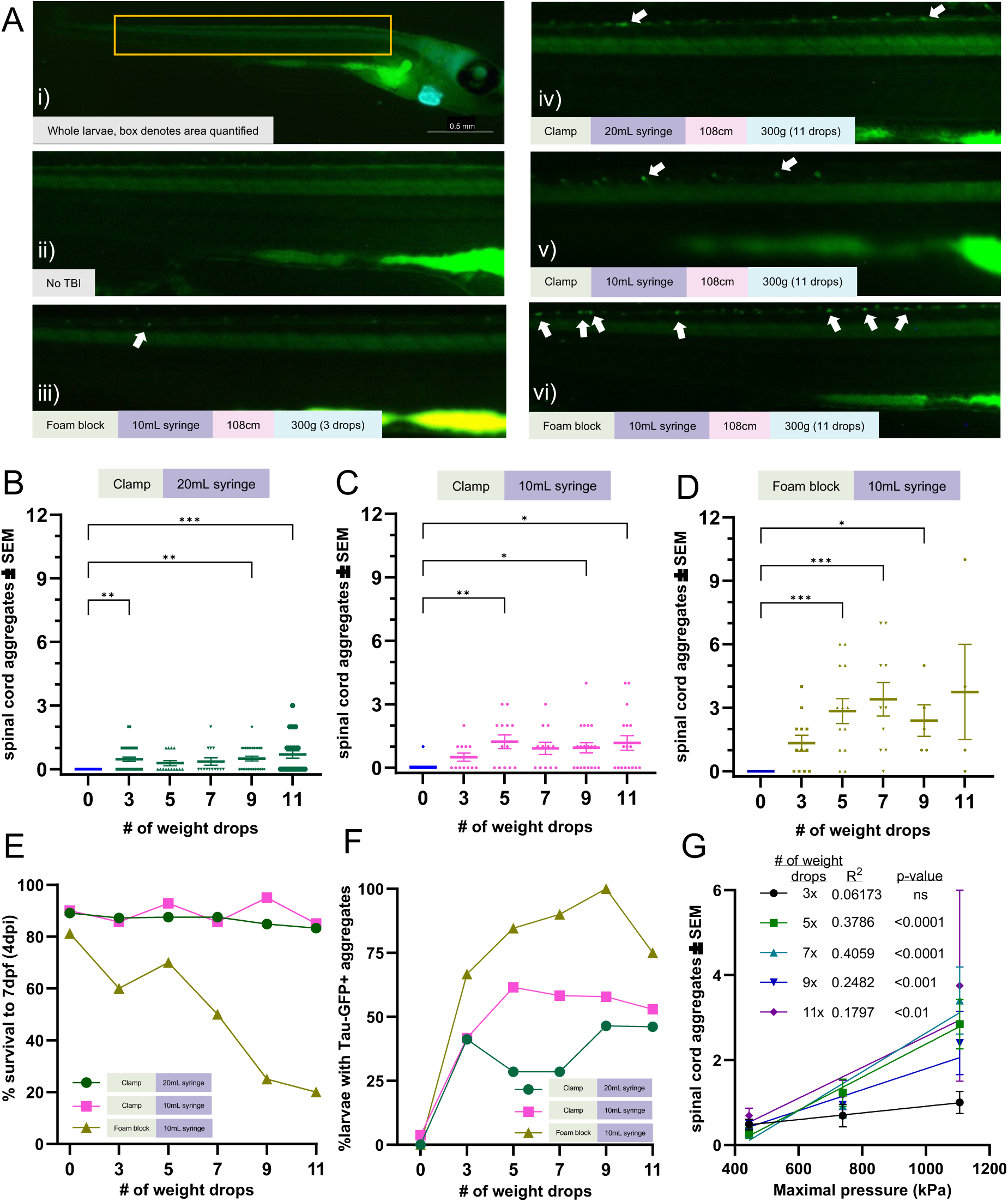
Tau aggregation increases with injury levels. Tau biosensor larvae were used to quantify how increasing injury intensity impacts the amount of detectable tau pathology. These transgenic larvae express full-length human tau and the human tau 4R domain linked to GFP, allowing for the visualization of tau aggregates as GFP+ puncta. Data represents individual larvae and bars represent +/- SEM and each experimental group was replicated at least twice. Representative images of larval zebrafish spines, where quantifiable GFP+ Tau puncta increase as the severity of injury and number of weight drops increase. Image i) depicts the full body of a larva for orientation and scale (0.5mm bar). White arrows denote GFP+ Tau puncta. Quantification of GFP+ tau aggregates in larvae injured in a 20mL syringe held by a clamp (N = 41, 34, 14, 14, 26, and 24 for the 0, 3, 5, 7, 9 and 11 drops respectively). **C**) Quantification of GFP+ tau aggregates in larvae injured in a 10mL syringe held by a clamp (N = 27, 12, 13, 12, 19, and 18 for the 0, 3, 5, 7, 9 and 11 drops respectively). **D**) Quantification of GFP+ tau aggregates in larvae injured in a 10mL syringe held by a foam block (N = 13, 12, 13, 10, 5, and 4 for the 0, 3, 5, 7, 9 and 11 drops respectively). **E**) Percent of zebrafish larvae that survived to 7 dpf after TBI using each TBI setup (N = equivalent to B-D). **F**) Percent of zebrafish larvae with one or more tau GFP+ puncta in each TBI model setup. **G**) Linear regression of mean spinal cord tau puncta (data from Fig.5 B-D) as a function of maximal pressure for larval injury groups that received differing numbers of weight drops. Every weight drop group except the 3 weight drop group significantly trended with maximal pressure. The 5 and 7 weight drop groups had the highest R^2^.

The lowest maximal pressure 20mL syringe held by clamp setup resulted in the least amount of tauopathy, with mean tau puncta counts and the percentage of larvae with tau remaining stagnant across weight drop groups (Fig. 5A(iv), 5B, p<0.01-0.001 vs control, 5F (green)). Larvae injured in the higher pressure 10mL syringe/clamp setup had a 1.7-to 4.31-fold increase in tauopathy versus the 20mL syringe/clamp group (Fig. 5A(v), 5C). 5 or more weight drop events increased detectable tau and the percentage of larvae with tau versus the 3 weight drop group (Fig. 5C, p<0.05-0.01 vs control, Fig. 5F (pink)). The highest pressure 10mL syringe/foam block TBI setup resulted in a 2.83 -9.96- and 2.31-3.71-fold increase in tauopathy versus the 20mL and 10mL syringe held by clamp groups, respectively (Fig. 5A(iii), 5A(vi), 5D). The mean tau count and percentage of larvae with tau trended with increased weight drops but decreased at the highest weight drop levels (Fig. 5D, p<0.001 vs control, Fig. 5F (brown)). The decrease in the percentage of tau GFP+ larvae, mean puncta counts and high variability for 9 and 11 weight drop groups is most likely because of high lethality (Fig. 5E). Increasing weight drop events can steeply increases mortality at higher pressure levels, therefore slightly lower injury levels may be best to optimize detectable tauopathy.

Multiple logistic regression analysis of these data predicted that each weight drop increased the odds of larvae having tauopathy by 9.9% and each 1kPa increase in maximal pressure increased the odds by 0.4%, or 49% per 100kPa (OR per hit: 1.099, OR per 1kPa: 1.004, Table 1). Linear regression showed tauopathy in the 5 and 7 weight drop groups scaled with maximal pressure the most (Fig.5G), suggesting 5-7 weight-drop events may be optimal, especially when considering optimal survival rates (Fig.5E). These results show that higher injury intensities and more injury events increase tauopathy in larval zebrafish, much like what is reported in humans, further validating the relevance of our assay.

## Discussion

In this study, we characterized the pressure dynamics in our larval TBI model to optimize model settings for the study of moderate-severe TBI. Modified TBI parameters like syringe size, drop weight mass, drop weight height, and syringe holder rigidity, all alter the blast pressure and injury produced (Figure 1). TBI pathophysiology is heterogenous between and within injury severity groups, yet current TBI classifications used in research treat TBI severity groups as a singular disease processes^67^. For example, most mild TBI patients often recover completely after injury but 30% of patients suffer from persistent debilitating pathology^68^, which underlines differences in pathophysiological progression and outcome for TBI patients. Studying multiple injury severities of TBI is important for the successful translation of preclinical research to clinically significant findings^69^. Our data shows that altering parameters of our TBI assay reliably changed the blast pressure created which in turn correlated with increased injury levels as shown by the markers we tested.

Our study addresses gaps in available TBI models in two primary ways: 1) Our model can be used as a high-throughput vertebrate assay with hundreds to thousands of zebrafish larvae undergoing TBI in a day. The number of individual specimens that can be studied increases the amount of heterogenous TBI pathology one would encounter in comparison to other animal models of TBI. Additionally, the high throughput nature of our TBI assay makes it an excellent system to easily test large numbers of potential clinically relevant pharmaceuticals. 2) Our model has bioethical advantages functioning as a clinically relevant Replacement^51^ alternative to higher sentience vertebrate models. Reduction strategies are also available with our model where pressure measurements can be used for pre-experiment optimization without the need for trial-and-error use of animals.

### Novel insights of the larval zebrafish TBI model

Our results display a strong relationship between impact intensity and multiple injury phenotypes. The ability to reliably create a range of injury phenotypes through altering injury intensity strengthens our model’s ability for studying heterogenous TBI outcomes. For example, our model could be used to establish a variety of injury levels and post-TBI phenotypes followed by transcriptomic/proteomic analysis of thousands of individual larvae to better understand pleiotropic mechanisms of TBI pathophysiology and outcomes.

We have also revealed that this TBI method introduces a non-linearity that results from changes in weight bounce, weight fall, the wobble of the syringe stand etc., suggesting that small forces can have an impact on injury. Careful experimental and assay design is needed when creating such TBI assays to ensure the similarity of injury levels. Furthermore, studies into the mechanical events behind injury induction may provide new information about how variability in injury events can impact the biological manifestation of injury phenotypes. We additionally found a biphasic relationship between injury intensity and locomotor seizure phenotypes. These results suggest that high-throughput examination of behavioural seizures require AI to detect severe seizures as injury increases, meaning that further refinement of available tracking systems can expand the high-throughput capabilities of PTS research in larval zebrafish.

### Optimal TBI settings for the study of moderate/severe TBI

The optimal dosing of injury within an experiment is attained by using the foam block and 10mL syringe with a large drop height. These settings combine to provide the broadest range of TBI pressures (spanning 33-1105kPa maximal pressure) and injury severity to the larvae by altering the mass of the weight dropped. Additionally, we suggest adjusting the guide tube diameter to match the diameter of the weight being used to reduce trial variability. If larval injury needs to be increased further than the maximum injury level obtained using the highest weight in the TBI setup, the number of weight drops per injury group can be increased to exacerbate injury, but researchers should be cognizant of the steep decline in larval survival^51^ (Fig. 5E). If the goal of the researcher is to establish an injury “dose-response”, maximal pressure measurements are an appropriate metric when modulating pressure through changing an individual parameter within the same TBI setup. When multiple TBI settings are changed at once the pressure dynamics created within the system can change in multiple ways (propensity for second pressure waves, time between weight bounces on the syringe plunger etc.) meaning the maximal pressure values <u>alone</u> may not be the best comparison.

### Characteristics of moderate-severe TBI in the larval zebrafish TBI model

TBI is categorized as mild, moderate, and severe based on criteria such as structural imaging, loss-of-consciousness (LOC), amnesia, and Glasgow Coma Scale score^70,71^. Some of these criteria cannot be measured in larval zebrafish but other metrics can be used to understand TBI severity at different pressure levels. In our injury groups, we saw that larvae experienced a stunned phenotype, seizures, and dementia pathology.

After injury, larvae are prone to falling on their side for long periods (up to hours for intense injury) and exhibit a suppressed response to stimuli, suggesting compromised neuronal function. Such injury phenotypes may be akin to a larval zebrafish LOC-like state that increases with injury severity, similar to how LOC increases with injury level in humans^71^. Further studies are needed to characterize this stunned phenotype as injury increases, but at this stage, the stunned phenotype is important to consider when designing experiments (especially behavioural) and could differentiate TBI severity in larval zebrafish.

PTS occurs in 5-7% of patients hospitalized with TBI, with injury severity being a primary risk factor, increasing to 11% and 35-50% for severe non-penetrating and severe penetrating TBI respectively^72-74^. Our larval zebrafish showed no seizure activity for low-intensity injuries, increasing injury further caused low-grade seizure responses like hyperactivity and whirlpool swimming, and in the most severely injured larvae, intense convulsive seizures were detected. The number of larvae experiencing seizure phenotypes also increased with the injury severity. Locomotor activity decreased with higher injury intensities, as reported in other TBI studies^21,63^, leading to decreased seizure detection via behavioural software. A combination of manual seizure scoring, calcium imaging^10^, and EEG assays may be of high value when detecting PTS after severe injuries in larval zebrafish.

PTS models are essential for developing pharmaceutical interventions for PTS and post-traumatic epilepsy (PTE). Acute PTS (seizure(s) <7 days post-TBI) are effectively treated with anti-seizure drugs and can often be prevented with early prophylactic intervention, whereas PTE (1 or more seizure >7 days post-TBI) cannot be prevented through prophylactic anti-seizure medication and 30-40% of epilepsy patients experience seizures that are drug resistant^75^. We have previously demonstrated that anti-seizure medications are successful in treating/preventing pathology following TBI in our larval zebrafish model^10^. Our zebrafish model’s similarity to the clinical etiology of TBI pathophysiology and high throughput potential make our paradigm an enticing option for dissecting the mechanisms of pharmaceutical-resistant PTS/PTE and testing potential anti-epileptogenic drugs.

TBI is an important factor of dementia as it is the primary cause of chronic traumatic encephalopathy (CTE) and a major risk factor Alzheimer’s disease (AD) ^76-80^. Repetitive mild TBI leads to tau pathology over years-decades in patients with CTE, but single moderate-severe TBI has also been linked to increased tau pathology^81,82^. Using our “tau biosensor larvae”, which express full-length human tau and Tau4R-GFP^10^, we measured tau pathology after varying TBI intensities to optimize detectable tauopathy. Low-pressure injuries resulted in little tauopathy, and high-pressure level injuries resulted in an abundance of tauopathy, displaying a trend that depended on injury severity. Our previous work quantifying tauopathy in this fashion (focused on the spinal cord as a convenient proxy for tauopathy in the brain) allowed us to identify and characterize various drugs and mechanisms regarding post-traumatic epilepsy^10^. Additionally, here we found that applying additional injury events increased the detectable tauopathy which is true in humans, further strengthening the validity of our model.

Optimizing the detection of tauopathy in our TBI model is important because of the potential to dissect pathomechanisms using our transgenic biosensor larvae. We have previously shown that PTS are a druggable link to tauopathy using these tau biosensor larvae in our TBI model, where blocking PTS with pharmaceuticals reduced dementia pathology^10^. Our recent review details current research connecting TBI to PTS and seizures to dementia, but a prominent gap exists in the literature since very few studies have directly tested the connection between TBI, PTS, and dementia^52^. The refinements discussed here in TBI assay settings to detect tauopathy and PTS provides us and others with easier access to a model that can further define the mechanisms between TBI, PTS, and dementias, providing further insight into potential therapeutics.

## Conclusion

Novel TBI models that reflect the heterogenous nature of TBI are imperative to understand the pathomechanisms which shape long-term clinical outcomes like post-concussion syndrome, PTS/PTE, and dementias. The larval zebrafish TBI method can be accurately optimized to reflect clinically relevant pathology and phenotypes of TBI. The stunned unresponsive phenotype, PTS, and tauopathy all increase with injury severity. This TBI method provides advantages that complement available TBI methods such as its high-throughput capabilities where potential therapeutics can be tested towards an array of different pathological events. Large numbers of larvae can be injured at once (hundreds to thousands in a day) increasing the scale of individual specimen injury that can be examined. These aspects of the TBI model make it an excellent candidate for the preclinical stage of drug discovery. Lastly, the larval zebrafish TBI model has bioethical Replacement and Reduction advantages. Since larval zebrafish are less sentient than other vertebrae models and initial injury settings can be optimized via an inexpensive pressure transducer without larvae present, reducing the number of animals injured.

## Author contribution statement

LFL designed experiments, collected data, created figures, and wrote the manuscript. TG collected data, contributed to writing, and provided feedback for editing. AHB provided intellectual ideas for experiment design and provided feedback for editing. SAWT provided data, intellectual ideas for experiment design, and provided feedback for editing. HA provided intellectual ideas for experiment design and provided feedback for editing. EAB provided intellectual ideas for experimental design, supervision, and feedback for editing. WTA provided intellectual ideas towards experimental design, supervision, and assisted in writing and editing. All authors approved the final version of the manuscript for submission.

